# A dynamic kissing model for enhancer-promoter communication on the surface of transcriptional condensate

**DOI:** 10.1101/2022.03.03.482814

**Authors:** Qing Zhang, Hualin Shi, Zhihua Zhang

## Abstract

Enhancers are key DNA elements that regulate gene transcription by activating promoters in eukaryotes. Many potential mechanistic models have been proposed to account for enhancer-promoter communication, including the formation of the condensed enhancer-promoter clusterings that may result from liquid-liquid phase separation. However, the detailed kinetic mechanisms by which condensates may mediate enhancer-promoter contacts remain unclear. Here, we characterized enhancer-promoter communication in condensates using a self-avoiding polymer chain model under several possible spatial constraints. When the chain was confined to the spherical surface or interior, we discovered a distinct relationship between average end-to-end distance and contour length, as well as a distinct distribution of end-to-end distance. Experimental molecular tracking data showed strong preference in support of the spherical surface confinement hypothesis. Based on such observation, we proposed a novel enhancer-promoter communication model in which enhancers dynamically “kiss” promoters along the surface of transcriptional condensate. The theoretical results may guide future experiments, as researchers continue to elucidate the regulation dynamics of gene expression.

## I. INTRODUCTION

In eukaryote, most genes can be regulated by distal cis-regulatory DNA elements called “enhancers” which have a major effect on cell phenotypes and fates [1]. Enhancers target promoters which are upstream regulatory elements of genes, the point at which transcription is initiated. However, enhancers may be separated from their targeted promoters by hundreds of KB to several MB of distance in the genome [2], or even located in different chromosomes [3]. Although critical, how enhancers and promoters find and communicate with each other remains largely unknown. Canonically, both stable and dynamic kissing models have been proposed [4]. In the stable model, enhancer-promoter (E-P) contacts are mediated by trans-factors, resulting in the formation of a relatively stable E-P complex. In the dynamic kissing model, E-P contact is more transient. The two models are sufficiently distinct. For example, to explain the transcriptional bursting phenomenon, whereby transcription from DNA to RNA can occur in pulses [5, 6], the stable model implies that extra regulatory signals are required for activation of the promoter, whereas in the dynamic kissing model, transcriptional bursting can be regarded as a natural result of the transition between enhancer binding to and unbinding from promoter.

These two models are also not mutually exclusive. For instance, when E-P contact is mediated by the droplet like condensates consisting of coactivators and transcriptional factors (TFs) [7, 8], the interaction can be categorized as exhibiting both stable and dynamic kissing modes [4, 9, 10]. Recent single-molecule tracking data (e.g., [11–13]) showed that extreme proximity between enhancer and promoter is not necessary for communication to occur [4, 10], suggesting that enhancers and promoters may both remain in condensates for a relatively long period of time and still stay highly dynamic.

The interplay between genome and condensates has been investigated through both experimentation and simulation (e.g., [14–16]). Condensates pull targeted genomic loci together, but exclude nontargeted regions [14]. The viscoelastic chromatin network constrains the growth of condensates [15]. The mutual repulsion between condensates and the chromatin network can lead to a clear separation between them [16]. These facts may provide clues that can explain the phase separation model of E-P communication. The attractive interaction between components inside transcriptional condensates, such as transcription factors (TFs) and their specific DNA binding sites, including enhancers and promoters, and the repulsive interaction between condensates and other genomic regions may, together, determine the random movements of enhancers and promoters, as well as their communication. This integrative effect of attractive and repulsive interactions between transcriptional condensates and the E-P DNA region may lead to an intriguing mechanism whereby enhancers communicate with promoters on the surface of condensates. However, a detailed model that replicates the dynamism and regulation of E-P contacts on the surface of condensates remains to be elucidated.

Therefore, in this study, we used the self-avoiding walk (SAW) model, in which a sequence of moves does not repeatedly visit any point on a lattice, to characterize random motions of DNA fragments containing enhancers and promoters under predefined spatial constraints, mimicking confinement brought by condensates that mediate communication. We combined the Flory theory, whereby the equilibrium conformation of a real chain is determined by the trade-off of repulsive energy and entropy loss [17] and simulation with the “pivot” algorithm, a dynamic Monte Carlo method that generates SAWs [18]. This allow us to formulate the relationship between average end-to-end distance (EED) and contour length and the dependence of this relationship on both size of the confinement sphere and proportion of the confined chain beads. With the scaling theory, which captures the invariance of certain relationships as the system changes in scale, we also obtained a theoretical formula to estimate the probability distribution of EED, which is comparable to experimental data. Comparisons between theoretical and experimental distributions of E-P distance exclude the mechanism of E-P random walk and communication within transcriptional condensate. The comparisons also indicate that condensate in large size and a high proportion of confined sites are needed for E-P random walking and communicating on the surface of transcriptional condensates. As an application of the condensate surface-communication model, we also simulated transcription bursting and explained the weak correlation between EP distance and gene expression observed in experiments. In conclusion, we proposed an E-P dynamic kissing model on condensate surface, which can be tested by more well-designed experiments.

## II. METHODS

### A. The SAW model for a polymer chain

To simulate the random walk of a DNA fragment containing enhancer and promoter, we considered a linear polymer chain composed of *N* identical hard spheres, which we hereinafter term as “beads” as distinct from condensate spheres, and numbered 1, 2, …, *N* sequentially. Two number-adjacent beads are always assumed to make direct contact. Thus, the distance between the centers of two number-adjacent beads equals bead diameter (*l*). The conformation of the chain can vary randomly, while still maintaining direct contact between two number-adjacent beads and avoiding overlapping between beads. Denote the coordinate of bead *i* as (*x_i_, y_i_, z_i_*). The contour length between bead *i* and *j* (*i* < *j*) is (*j* – *i*) * *l*. The (internal) EED for the subchain consisting of beads *i*, .., *j* is the 3-D Euclidean distance between beads *i* and *j* (*i* < *j*), i.e., 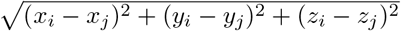. In both theory and simulation, we derived the average EED as a function of contour length and the probability distribution of EED.

### B. Simulation

We adopted a Monte Carlo method, a variant of the “pivot” algorithm [18], to simulate the random walk of the self-avoiding polymer chain in different spaces. The initial conformation of the chain was typically generated by randomly adding beads, one by one, from one end to the other end. If the newly added bead in producing the original conformation violates spatial constraints, it will return to an earlier step which is arbitrarily or randomly chosen. The conformation of the chain was adjusted by choosing a “pivot” bead randomly and doing a space-dependent random operation on a part of the chain. A series of stochastic operations produced an ensemble of conformations, from which the statistics were made, including the average and distribution of EED. The pivot algorithm is ergodic [18], an observation supported by the consistency between the time-average and ensemble-average results. Instead of simulating on lattices [18], we followed the coordinate of each bead at each step in 3-D continuous space. Specific details of the SAW simulation with the pivot algorithm in the constraint spaces of interest are presented below.

- *In 3-D space*. A bead is randomly chosen as a “pivot”, and the beads on both sides are defined as left and right arms, respectively. Using this pivot bead as the origin, rotate the shorter arm centered on the pivot bead. The rotation has two random orientation angles, *θ* (∈ [0, *π*)) and *ϕ* (∈ [0, 2*π*)). If the rotated arm overlaps with the other arm, the rotation is rejected, and the count number of last conformation is increased by ‘1’. The rotation operations are performed at least 2 × 10^5^ times. Gyration radius and EED are calculated for each step. The simulation does not stop until these two variables show no increasing or decreasing trend for a long enough time.
- *In 2-D space (i.e., plane*). The rotation operation has only one random orientation angle, *ϕ* (∈ [0, 2*π*)). The axis of rotation is perpendicular to the walking plane and passes through the center of the pivot bead.
- *On a spherical surface*. The procedure is similar to that for 3-D space; however, the rotation has only one random orientation angle, *ϕ* (∈ [0, 2*π*)). The rotation axis passes through centers of the pivot bead and the sphere.
- *Inside a sphere*. The procedure is, again, the same as that for 3-D space, except that the rotation is also rejected when one bead in the rotated arm walks out of the sphere.
- *A certain proportion of beads confined to a spherical surface*. Define the left and right arms the same as that for 3-D space. Define the beads between two adjacent confined beads as one subchain. Randomly choose one “pivot” bead. Define the subchain where the pivot bead exists as the pivot subchain. Define the left (or right) part of the pivot subchain as the left (or right) local arm. If the pivot bead is confined to the surface, rotate the shorter arm with a random orientation angle (*ϕ* ∈ [0, 2*π*)) around the axis passing through centers of the pivot bead and the sphere, just as the rotation when the whole chain is confined to the spherical surface. If the pivot bead is not confined to the surface, rotate the right (or left) local arm as in the operation in 3-D space. Then rotate the whole pivot subchain to make its right (or left) end return to the surface around the rotation axis passing through the left (or right) end and perpendicular to the plane containing the pivot bead and the two ends of the pivot subchain. Finally, rotate the right (or left) subchains of the pivot subchain to reconnect with the adjusted right end of the pivot subchain around the axis passing through the spherical center and perpendicular to the plane containing the spherical center and the unadjusted and adjusted right (or left) end of the pivot subchain.
- *A certain proportion of beads confined inside a sphere*. The procedure is the same as that for 3-D space, except that the rotation is also rejected when one confined bead leaves the sphere.

Unless otherwise specified, we simulated random walks of a chain with 401 beads. The free ends behave a little differently compared to internal ends, the latter being more reflective of reality. To exclude the free end effect, 300 internal beads were considered to compute the contour length and EED. The relationship curves of EED and contour length from theory and simulation were both fitted with the power-law formula *R* ∝ *N^ν^*. The fitted exponent *v* (named “power-law” exponent) was used to reflect the spatial degree of freedom of the walking chain, to some extent. The probability density function (PDF) of simulated EED was scaled with the average, i.e., the EED in each conformation was divided by the average over the time.

### C. Flory theory and scaling method

The conformation of a self-avoiding chain is controlled by the balance between two free energy sources: the repulsion energy between beads in the chain and the entropy loss owing to deviation from the conformation of an ideal chain [17]. The former contributes to the swelling of the chain, while the latter limits the swelling. Flory theory is a simple model for phenomenologically capturing this balance [17]. As an example, below we show how the Flory theory works when a self-avoiding chain walks in 3-D space.

Denote the number of beads in the chain as *N*, the diameter as *l* (assumed to be 1 for simplicity), and the instantaneous EED as *r*. Use *R* to denote the average EED 〈*r*〉, which reflects the shrunken size of the chain. Under the mean-field theory, chain beads can be seen to distribute uniformly in a sphere with diameter *R*, and thus the density of beads is 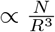. The interaction energy for one bead with others is 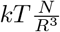 approximately. Then the interaction energy contained in the chain is around 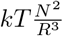. The energy contributed by entropy can be approximated as 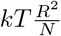 [17]. So, the total free energy can be represented by

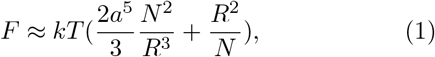

where *a* is a scaling constant. When the total free energy reaches the minimum, one can derive that

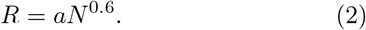

When the chain walks in 2-D space, the total free energy can be derived in a similar way as

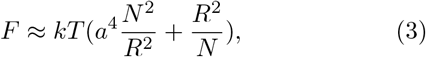

where *a* is a scaling constant, and the minimization of the total free energy leads to

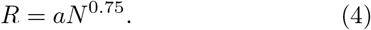

Equations 2 and 4 indicate that the average EED is a power-law function of the contour length of the chain with the exponents of 0.6 and 0.75 in 3-D and 2-D space, respectively.

We scaled the simulated or experimental EED by dividing each instantaneous distance by the average 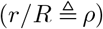. In the elaborate theory ([19–21]), the probability density function of such a scaled EED for a long SAW chain is derived as

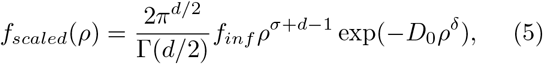

where *d* is the dimensionality. Then the probability density function of the original EED can be estimated by

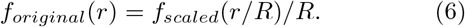

Based on the Flory theory and scaling method, the average, scaled and original distributions of the EED are quantitatively expressed as Eqs.2 and 4–6, respectively. The SAW simulation well reproduced these conformation features of self-avoiding polymer chain in 3-D and 2-D Euclidean spaces [Appendix A and Fig. 7].

## III. RESULTS

### A. The entire polymer chain walking on the surface of a sphere

A chromatin network can constrain condensates, even while condensates can expel chromatin [14–16], which may lead to random walk and communication of enhancer and promoter on the surface of transcriptional condensate. First, consider an extreme case in which all beads in the polymer chain walk on the surface of a sphere.

The EED on the spherical surface (*R_SS_*) can be derived in a manner similar to when the chain walks in 2-D space. The free energy of the chain walking on the spherical surface can be written as

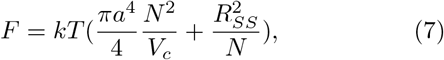

where the volume occupied by the chain can be estimated by 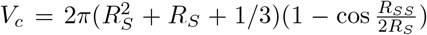. The steady state *R_SS_* can be derived from 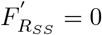, i.e.,

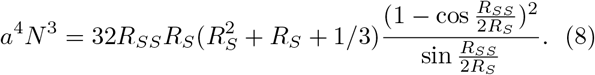

As shown by Figure 1 (a), the EED in 3-D space can be related to *R_SS_* based on the geometrical relationship

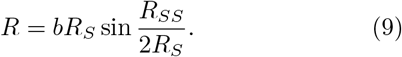

**FIG. 1.**
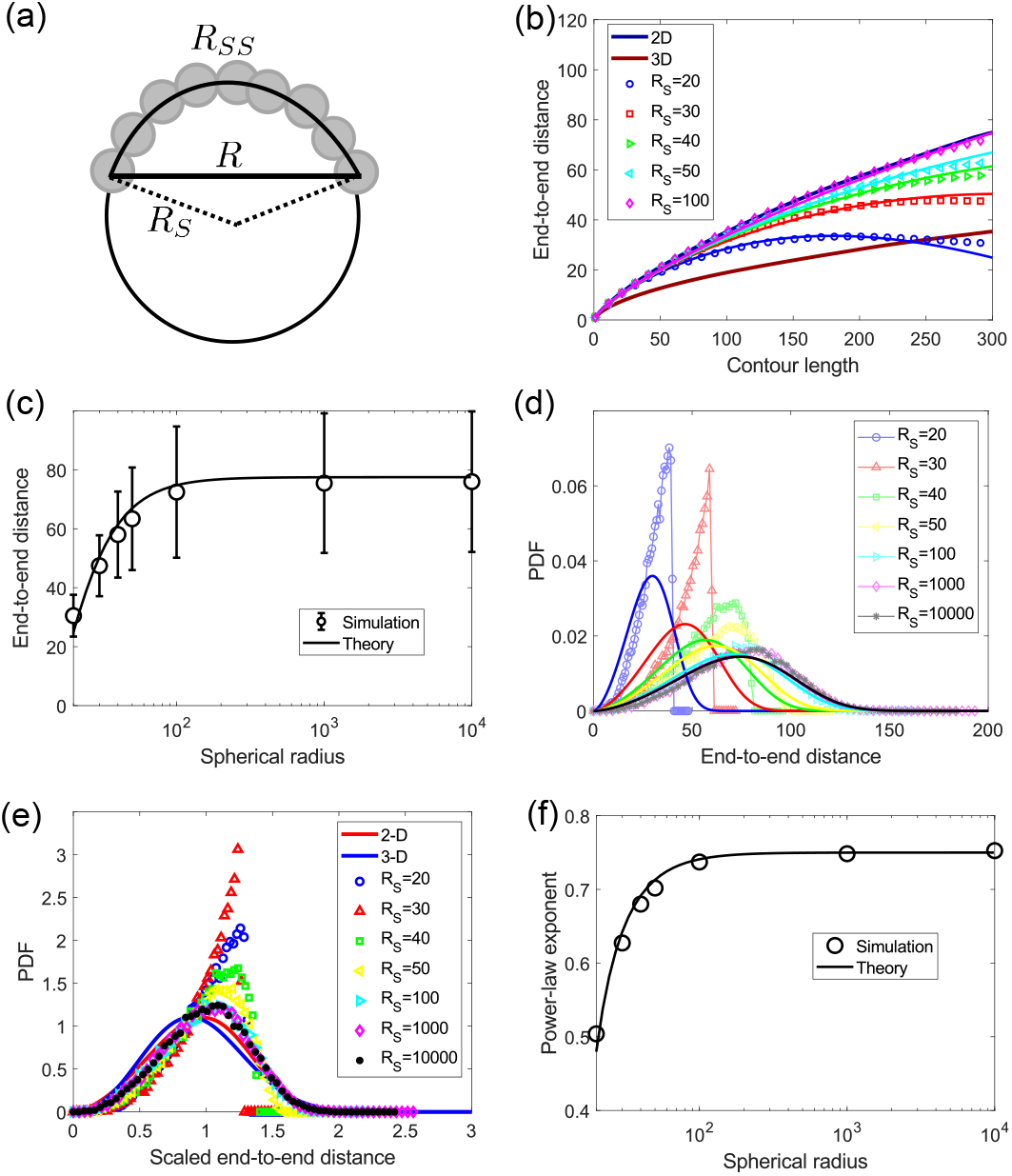
Spherical size affects random walk of DNA chain which is entirely confined on the surface of the sphere. (a) The schematic diagram. *R_S_* denotes the spherical radius. *R* and *R_SS_* denote the EED in 3-D space and on the spherical surface, respectively. (b) The relationship between EED and contour length is changed with *R_S_*. The result with a large sphere (e.g., *R_S_*=100) is similar to that in 2D space. When spherical size is limited, the relationship curve is lowered. Symbols (different *R_S_* values) and thick lines (2D and 3D) denote results from simulation; other lines denote theoretical results obtained by fitting simulation results with Eq. 10. Parameters: *a* = 1.28 and *b* = 1.68. (c) EED increases gradually with *R_S_* and enters a plateau at *R_S_* ~ 100. Error bars denote standard deviation. (d) The probability distribution of EED. (e) The scaled probability distributions of EED on small spheres do not collapse to theoretical 2D and 3D distributions. (f) Power-law exponent increases gradually with *R_S_* and enters a plateau (~ 0.75) at *R_S_* ~ 100.

Equations 8 and 9 give the dependence of average EED on contour length and spherical radius. When *R_S_* ≫ *R_SS_*, Eq.8 decays to the 2-D space form, i.e., *R_SS_* = *aN*^0.75^, and Eq. 9 is simplified to

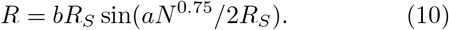

Below we use this simplified formula instead of Eqs. 8 and 9 to fit the simulation results since the difference is small.

The simulation shows that the average EED decreases with decreasing spherical radius [Fig.1 (b) and (c)]. The relationship between the average EED and contour length deviates from the 2-D power-law form, as shown by Eq.4, but gradually as the spherical diameter decreases [Fig.1 (b)]. With Eq. 10, the theory well captures the dependence of EED on the radius [Fig.1 (b) and (c)]. The shape of EED distribution is also changed as the radius decreases (Fig.1 (d)). When the sphere is small enough, the EED distribution falls to zero sharply, around *R* = 2*R_S_*, which clearly indicates the effect of spherical size. By scaling with the average, the EED distribution can collapse to the curve given by Eq. 5, but only when the sphere is large enough [Fig.1 (e)]. Compare with the distribution of experimental E-P distance (see below). This, in turn, suggests that only a sufficiently large condensate sphere will allow E-P walking and communicating on a spherical surface. The exponent obtained by fitting the EED-contour length relationship with the power function (*R* ∝ *N^ν^*) also shows a decrease with decreased spherical radius [Fig.1 (f)]. This result suggests that the spatial degree of freedom of the chain is increased from that of 2-D space when the spherical size is reduced.

### B. The polymer chain is partially confined to the surface of a sphere

Enhancers and promoters could be confined to transcriptional condensates owing to the specific binding with TFs and other factors. Regions neighboring enhancer and promoter may also contain some pseudo binding sites with certain strengths. The number of these binding sites affects the formation of phase-separated condensates [22], and it may also affect the walking pattern of enhancers and promoters. In this section, we study the effect of a change in the proportion of chain beads confined to the spherical surface, given that confined beads are evenly distributed in the chain.

Assume that the spherical diameter is sufficiently large so that the spherical surface, i.e., walking space of confined beads, could be considered as 2-D space (see above results), while the beads far from the confined beads could be described as walking in 3-D space [Fig. 2 (a)]. Denote the proportion of confined beads as *ϕ*. In principle, the confinement on the limited beads also affects the walking of unconfined beads, and the effect diminishes according to the distance of the beads to their closest confined sites increases. That is, the farther away from the confined sites, the less restricted the beads are and the higher the freedom of walking. Thus, the neighbors of confined beads should walk in a manner that is close to walking in 2-D space, but the beads farther away from the confined beads are more likely to walk in 3-D space. Applying the Flory theory, in this case, the free energy contributed by entropy is still proportional to *R*^2^/*N*, while the interaction energy should be a compromise between those interaction energies in 3-D and 2-D space. To construct the interaction energy, we partitioned the chain into two parts: (1) confined beads and their neighbors (denoting the proportion by *f* (*ϕ*)) and (2) the remaining beads (the proportion is 1 – *f*(*ϕ*)). For simplicity, we assume that beads in the first class all walk in 2-D space, whereas beads in the second class all walk in 3-D space. Thus, the interaction energy can be roughly expressed as a linear combination of those in 2-D and 3-D space and total free energy can be represented by

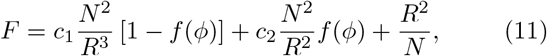

where *f*(*ϕ*) and constants *c*_1_ and *c*_2_ can be trained by the simulation. The linear form of *f*(*ϕ*) cannot lead to a good agreement with the simulation results, so we used a power form, i.e.,

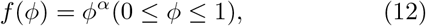

where the exponent *α*, as well as *c*_1_ and *c*_2_, can be determined by fitting with simulation results. When *ϕ* = 0, *f*(*ϕ*) = 0, and Eq. 11 decays to Eq.1, i.e., free energy in 3-D space. When *ϕ* =1, *f*(*ϕ*) = 1, and Eq. 11 decays to Eq.3, i.e., free energy in 2-D space. These observations indicate that the theory works in the two extreme cases of all beads confined and no beads confined. To compare with the simulated average results, one can derive the average EED R at the steady state by solving 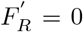, i.e.,

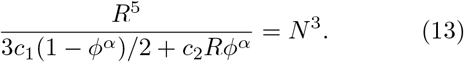

As shown in Figure 2 (b), the relationship between the average EED and contour length from the simulation changes from 3-D form to 2-D form with the proportion of confined beads in the chain increasing. Moreover, the EED becomes longer with the increasing confined proportion [Fig 2. (c)]. The theoretical relationship among average EED (R), confined proportion (*ϕ*), and bead number (*N*) given by Eq.13 fits the simulation well [Fig. 2 (b) and (c)]. EED increases with confined proportion because the confinement of beads increases self-avoiding between beads and then swelling of the chain. Like the average EED, the variability of EED also increases with the confined proportion [Fig. 2(c) and (d)]. Furthermore, the distribution of EED divided by the average collapses to 2-D and 3-D versions of Eq. 5 [Fig. 2(e)]. Accordingly, we theoretically estimated original distribution of EED by combining theoretical formulas for the average and the scaled probability distribution (i.e., Eqs. 5 and 13) (see Eq. 6), which is consistent with the simulated distribution [Fig. 2(d)].

**FIG. 2.**
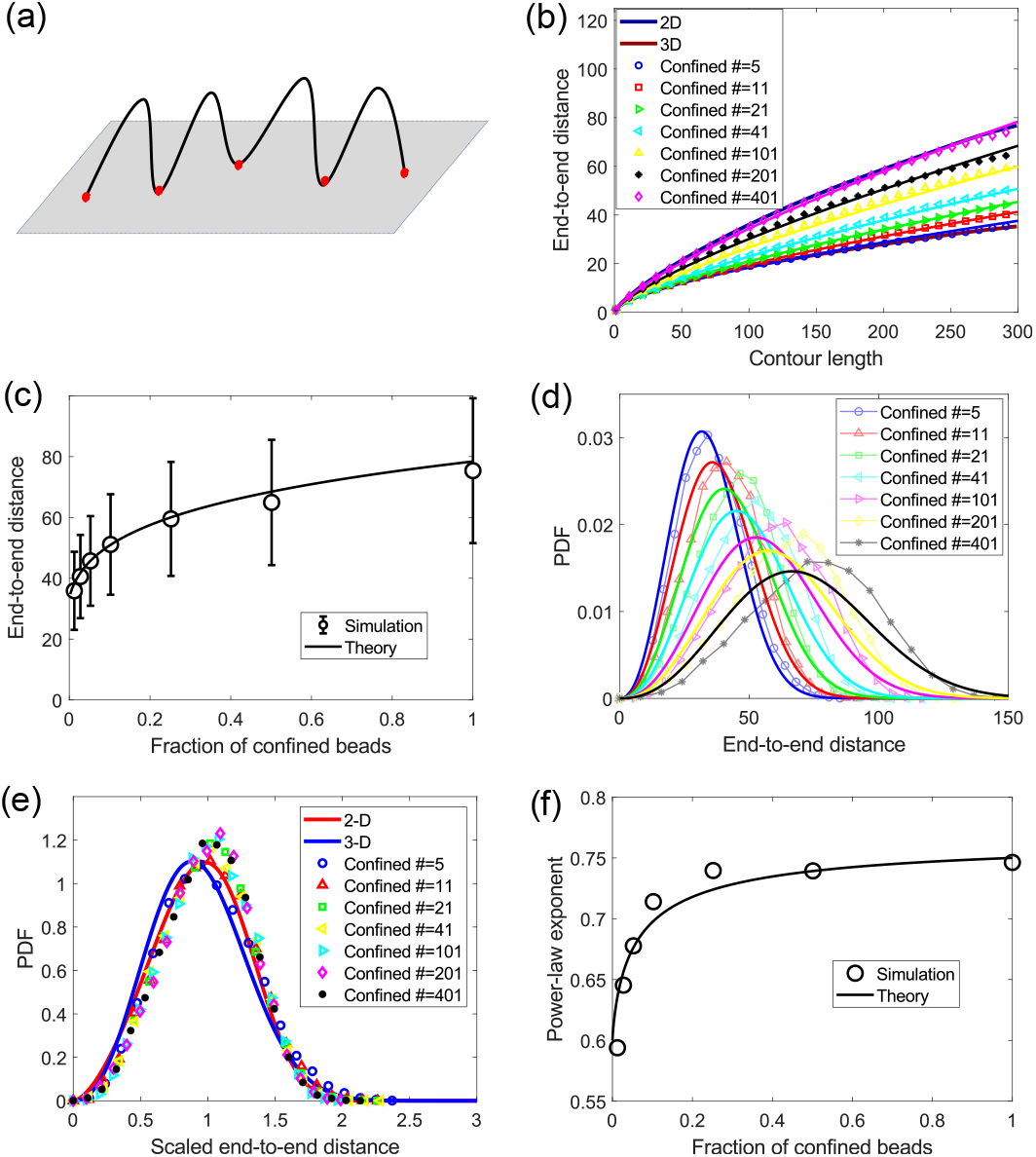
DNA chain randomly walks with some beads confined on the surface of a large sphere (*R_S_* = 10^4^). (a) The schematic diagram. Black line and red points represent the chain and confined sites, respectively. (b) The relationship between EED and contour length is changed with the number of confined beads. Few beads confined lead to the 3D result, whereas all beads confined lead to the 2D result. Symbols (different confined numbers) and thick lines (2D and 3D) denote results from simulation; other thick lines denote theoretical results obtained by fitting simulation results with Eq.13. Parameters: *α* = 0.8, *c*_1_ = 1.5, *c*_2_ = 1.4. (c) The EED increases with the proportion of confined beads. Error bars denote standard deviation. (d) The probability distribution of EED. (e) The scaled probability distributions of EED for different confined bead numbers all collapse to theoretical 2D and 3D distributions. (f) Power-law exponent increases with the proportion of confined beads from 0.6 (3D exponent) to 0.75 (2D exponent).

The results also show that EED and contour length has an approximate power-law relationship and that the exponent increases from ~ 0.6 (for 3D) to ~ 0.75 (for 2D) when the confinement proportion increases from 0 to 1 [Fig.2. (d)]. This result can be understood based on a scaling method. One can imagine the region between two adjacent confined beads as a higher-level bead, termed “meta-bead”. Then, the walk of the chain can be characterized by separately describing the walk of beads inside one meta-bead and that of meta-beads in the whole chain. We simulated the walk of beads within one meta-bead. The obtained EED (distance between two adjacent confined beads) as a function of contour length (the number of involved beads denoted by *g*) agrees with a power function with the exponent of 0.6 [Fig.8 (a)], indicating that the beads within the meta-bead walk in 3D space. We assume that the meta-beads walk in a space with a power exponent *v* which depends on *g*. Taken together, the EED of the entire chain can be represented by

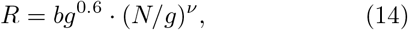

where *b* is constant. Learned from the simulation, the exponent *v* can be expressed as a function of *g* by

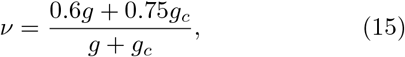

where *g_c_* is constant. Equation 15 shows that *v* increases with *g*; when *g/g_c_* ≈ 0, *v* = 0.6, while when *g* ≫ *g_c_, v* = 0.75. This is understandable because the effect of the confinement becomes less when the number of beads within one meta-bead (*g*) is bigger; meta-beads tend to be confined on the surface when *g* → 1, but tend to leave the surface when *g* → ∞. The decrease in the confinement effect can also be reflected by the increase in the relative standard deviation of EED [Fig.8 (b)]. The scaling method represented by Eq.14 also captures the effect of confinement on the relationship between EED and contour length and the power-law exponent [Fig.8 (c-d)].

If the size of the underlying transcriptional condensate is not large enough, then both effects of limited condensate size and limited confined sites need to be considered. The mixed effect can be described based on Eqs.9 and 13. Denote the solution of Eq.13 by *R_C_*, and substitute the approximate relation *R_SS_* ≈ *R_C_* into Eq.9, which yields

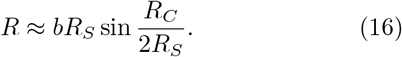

Equation 16 roughly captures the simulated relationship between average EED and contour length with different confined proportions and limited spherical size [Appendix Fig.9 (a)]. Appendix Fig.9 (b) and (c) present the dependence of average EED and power-law exponent on both spherical radius and confined proportion. These relationships could be examined by more specifically designed experiments.

### C. The polymer chain entirely or partially walking inside a sphere

To examine the scenario in which the enhancer communicates with the promoter inside the condensate, we tested the following constraints.

First, we consider the ideal situation in which all chain beads are confined to walk inside a sphere. Under the Flory theory, the free energy in this case can be derived by adding the interaction energy between the chain and the confining spherical surface (*F_SC_*) in the free energy for 3-D space, i.e.,

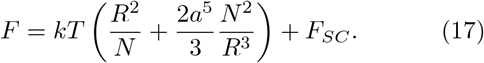

To derive *F_SC_*, we estimated the probability of the chain contacting with the confining spherical surface using the mean-field method. If the coordinate center of the chain is fixed, all possible conformations of the chain, on average, produce a spherically symmetric position distribution of chain beads, which we call “spherical chain cloud”. In the chain cloud, we defined a chain sphere as the space wrapped by the boundary that consists of points most likely contacting the restraining spherical surface and denoted by the radius *R_I_*, which reflects the minimal distance from the center of chain sphere to the surface of confining sphere [Fig.3 (a)]. The chain center is limited to movement just inside a spherically feasible space with the radius of *R_S_* – *R_I_*. Thus, the probability of the chain contacting the confining spherical surface (*p_contact_*) can be approximated by the probability of the chain center moving to the border of the feasible space, i.e., 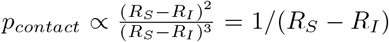. For simplicity, we assume *R_I_* = *cR*. Finally, *F_SC_* can be expressed as

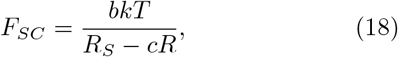

where *b* is a constant. The average EED of the chain at the steady state can be obtained by solving 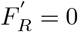, i.e.,

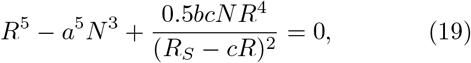

where *a* has the same value as in Eq. 2, and *b* and *c* can be determined by fitting simulation results. Interestingly, fitted c is around 1, indicating *R_I_* ≈ *R*. The effect of spherical size on the average EED is well captured by Eq.19 [Fig.3 (b) and (c)]. As shown by Figure 3 (b), the average EED changes with contour length more slowly than that for 3-D space when the spherical radius is smaller than a certain value (diameter threshold), while it is the same as that for 3-D space when the spherical radius is higher than the threshold. Correspondingly, Figure 3 (c) shows that the average EED does not change with the radius when the sphere is large enough, while it decreases with the decreasing radius when the radius is smaller than the threshold. Furthermore, the probability distribution of EED becomes narrower when the radius decreases under the threshold [Fig.3 (d)]. Irrespective of the radius, the scaled probability density distribution of EED collapses to theoretical curves for 2-D and 3-D space [Fig.3 (e)]. The power-law exponent obtained by fitting the simulated EED as a function of contour length decreases to become smaller than 0.6, corresponding to a higher spatial degree of freedom when the radius is smaller than the threshold (Fig.3 (f)). The differences between results shown by Figure 3 (b) and (f) and Figure 1 (b) and (f) could be used to distinguish whether the enhancer communicates with the promoter on the condensate or inside the condensate. For example, the comparison between the theoretical power-law exponents and the exponent inferred from experimental data (see below) more supports the surface-communication model.

**FIG. 3.**
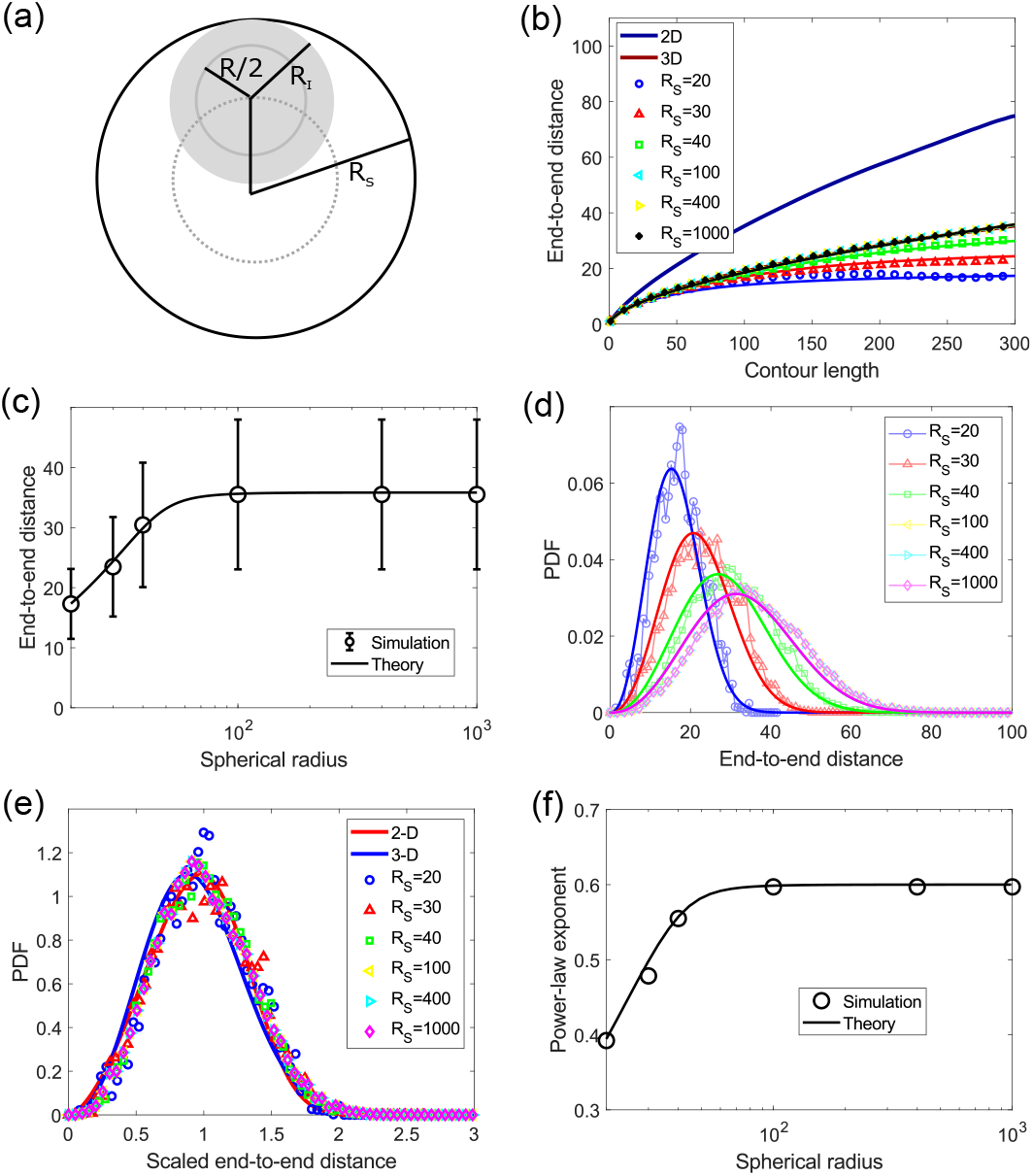
Effects of changing spherical size when the chain walks inside the sphere. (a) The schematic diagram. *R_S_* denotes spherical radius. *R* denotes the EED of the considered chain. *R_I_* denotes the distance of the chain center from the spherical surface on average. (b) The relationship between EED and contour length is changed with RS. The result with a large sphere (e.g., *R_S_* ≥ 100) is similar to that in 3D space. When sphere size is limited, the relationship curve is lowered. Symbols (different *R_S_* values) and thick lines (2D and 3D) denote results from simulation; other lines denote theoretical results obtained by fitting simulation results with Eq. 19. Parameters: *a* = 1.17, *b* = 30, and *c* =1. (c) EED increases gradually with *R_S_* and enters a plateau at *R_S_* ~ 50. Error bars denote standard deviation. (d) Probability distribution of EED. (e) The scaled probability distributions of EED for different spherical sizes all collapse to theoretical 2D and 3D distributions. (f) Power-law exponent increases gradually with *R_S_* and enters a plateau (~ 0.6) at *R_S_* ~ 50.

Second, we asked if only partial beads are confined to randomly walk inside the condensate sphere owing to specific binding with TFs or other condensate components, whereas others are allowed to walk both inside and outside of the condensate sphere. To test this mechanism, we still defined the proportion of confined beads as *ϕ* to see how EED changes as *ϕ* changes. When the sphere is large enough (*R_S_* ≫ *R*), results show that EED does not change with the proportion of confined beads (Fig.4 (a)). This is because the likelihood of the chain interacting with the surface of the large sphere is minimal, which means that the chain just walks in 3-D space, no matter how many beads are set to be confined. When the sphere is small (*R_S_* ~ *R*), as shown by Figure 4 (b) and (c), EED decreases with the confined proportion from the free 3-D case to the case of certain confined proportion (~ 10%), whereas it only changes a little when the confined proportion increases from ~ 10% to 100%. This observation suggests that partial confinement of the chain based on specific binding with internal components can lead to the chain entirely waking inside the small sphere.

**FIG. 4.**
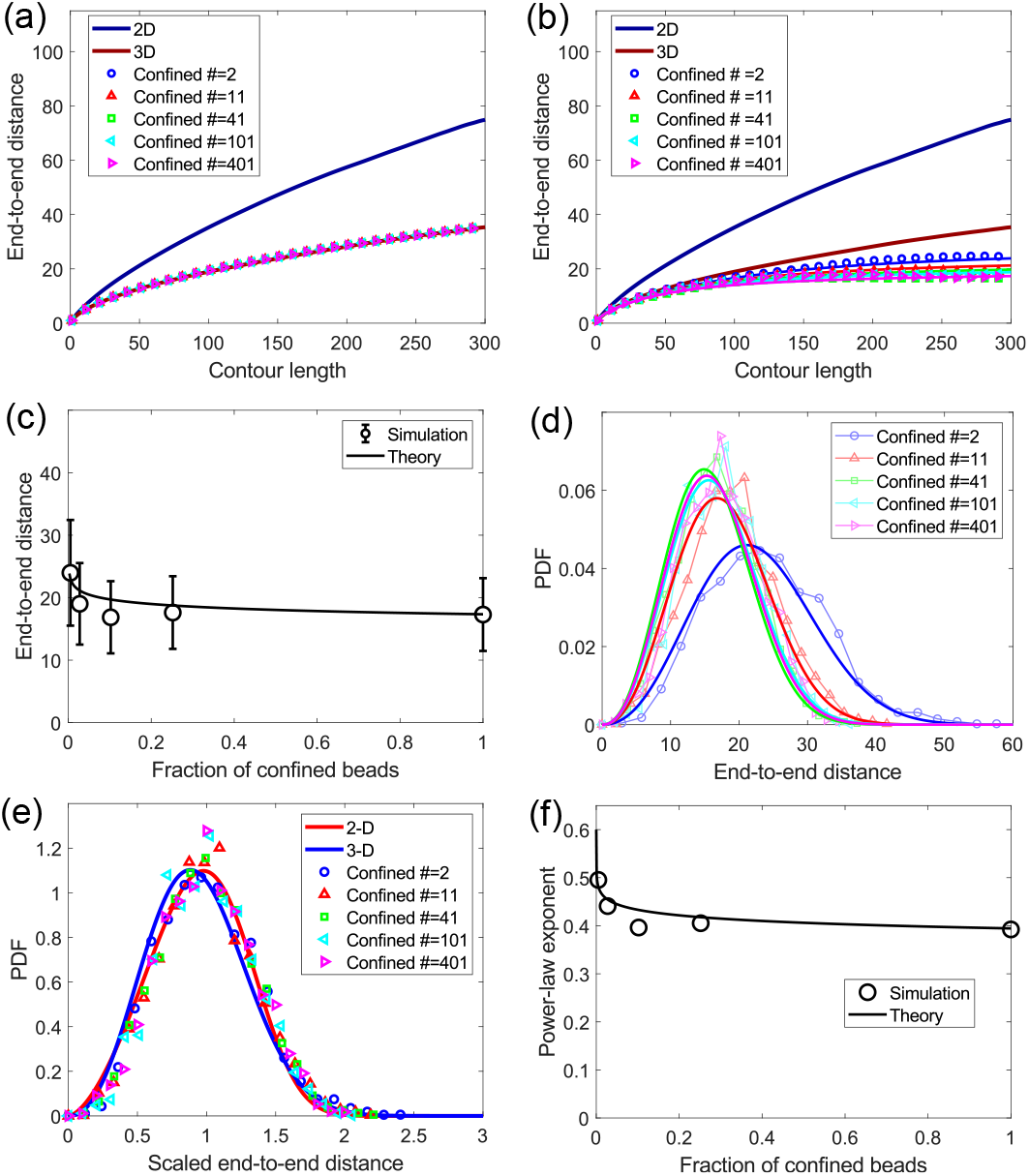
Random walks of the chain with certain proportions of beads limited inside a large or small sphere. (a) The relationship between EED and contour length is the same as that for 3D space, independent of confined bead number, when the confining sphere is large enough (here *R_S_* = 10^4^). (b-f) Results for a small confining sphere (*R_S_* = 20): (b) The relationship between EED and contour length is changed according to the number of confined beads. Symbols (different confined numbers) and thick lines (2D and 3D) denote results from simulation; other lines denote theoretical results obtained by fitting simulation results with Eq.21. Parameters: *a* = 1.17, *b* = 30, *c* =1, and *α* = 0.07. (c) EED decreases according to the proportion of confined beads when *ϕ* < 0.1, while it changes little when *ϕ* > 0.1. Error bars denote standard deviation. (d) The probability distribution of EED. (e) The scaled probability distributions of EED for different confined bead numbers all collapse to theoretical 2D and 3D distributions. (f) Power-law exponent decreases according to the proportion of confined beads from 0.6 (3D exponent).

The case of limited beads confined inside a small sphere can also be quantified theoretically by extending the theory for the full confined case. In the limited case, the interaction energy *F_SC_* represented by Eq.18 needs to be modified. When the confined proportion *ϕ* reduces, the feasible space of the chain center should become bigger, and the corresponding radius (*R_fs_*) should become larger than *R_S_* –*cR*. When only two ends are confined (even the contour is very long), the center of the chain, on average, can be close to the surface, but still be confined inside the sphere, i.e., *R_fs_* is still no larger than *R_S_*. So, we can approximate *R_fs_* to *R_S_* when the confined proportion *ϕ* is close to zero. Under this consideration, we can simply represent *R_fs_* by *R_S_* – *cϕ^α^R*, where the exponent *α* is a constant between 0 and 1. Then Eq.18 can be modified to

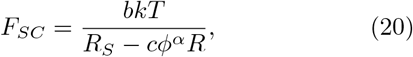

and Eq. 19 for deriving optimal *R* can be updated to

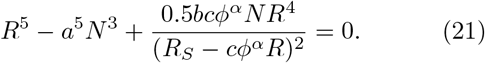

Figure 4 (b) and (c) show that Eqs. 20 and 21 can basically reproduce the decrease in EED with the confined proportion from the simulation. The variance of EED decreases with the confinement proportion in a manner similar to that of the average [Fig.4 (c) and (d)]. By dividing EED over its average, the probability distribution of the scaled EED under different confinement proportions collapses together [Fig.4 (e)] to the theoretical 2-D and 3-D scaled distributions as shown by Eq.5. The original probability distributions of EED estimated with Eqs. 5, 6 and 21 are in agreement with the simulated distributions [Fig. 4 (d)]. The power-law exponent also shows a decrease, along with the confinement proportion [Fig. 4 (f)], indicating a higher spatial degree of freedom than 3-D free space. The high spatial degree of freedom when the chain entirely or partially walks inside the condensate is contrary to the low spatial degree of freedom of the E-P region, as can be inferred from experimental data (see below), thus excluding the possibility that enhancers communicate with promoters inside transcriptional condensate.

### D. Single-molecule tracking data support a possible 2-D random walk model of E-P communication

Recent single-molecule tracking (e.g., [12, 13]) produced the trajectories of enhancer and promoter randomly walking and gene transcriptional bursts. With the data for all related trajectories in single cells, the distribution of E-P distance can be derived [Fig. 5 (a)]. The EED distribution of *in vitro* DNA fragments with different contour lengths has been observed to collapse to each other when the EED distribution is scaled by 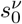 (*s*_0_ is contour length; *ν* is the power-law exponent in 3-D space) [23]. To test if the DNA region of interest with different E-P contour lengths always randomly walks in 3-D free space with three datasets from [12, 13], we scaled the distribution of E-P distance by dividing the distance by 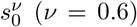, but the distributions did not collapse together [e.g., Fig. 5 (b)], indicating the existence of an alternative constraint space wherein the enhancer is walking along with promoter. When E-P distance is longer, communication will become more difficult, increasing the likelihood that the enhancer walks in 3-D space with the promoter. According to the above theoretical results, a stronger constraint leads to a larger scaling exponent. Figure 5 (b) shows that the shorter E-P contour corresponds to a longer scaled E-P distance (on average), which implies that the shorter contour tends to need a scaling exponent larger than 0.6 for the collapse. This is supported by the scaling results with the average considering the small difference in the scaling constant between different spaces: the distribution of EED divided by the average basically collapses to the theoretical 2-D and 3-D scaled distributions, irrespective of contour length [Fig. 5 (c)].

**FIG. 5.**
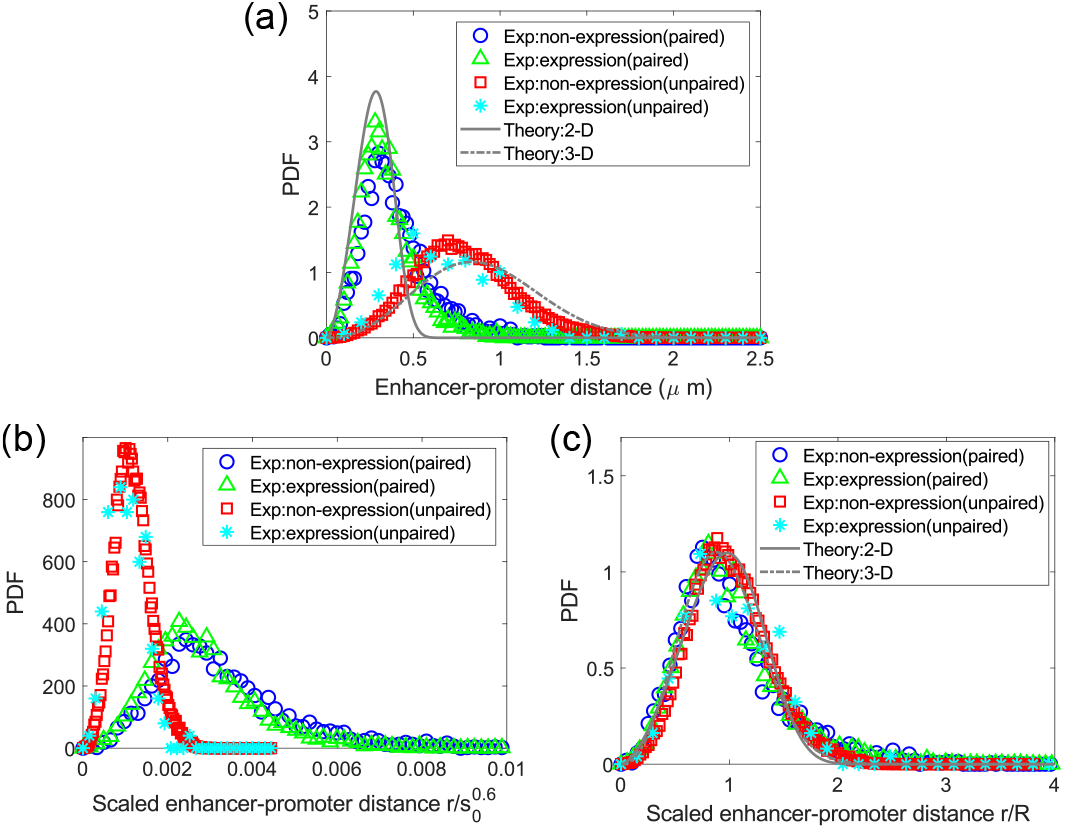
Probability distribution of E-P distance with different contour lengths is captured by the Flory theory and scaling method. The experimental data from Chen et al. [12] were used. Four datasets were considered based on the (homie) insulators as paired or unpaired and the gene as transcribing (expression) or not (non-expression). Related parameters are given in Appendix C2. (a) Probability density function (PDF) of original E-P distance was well estimated by the theory described by Eq. 24. (b) Probability density function (PDF) of scaled E-P distance cannot collapse together when the E-P distance is divided by 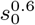 (*s*_0_ is the effective contour length). (c) Probability density function (PDF) of scaled E-P distance collapses to the theoretical 2-D and 3-D scaled distributions when the E-P distance is divided by its average.

Scaling with the average can be related to scaling with the power function by assuming a power-law relationship between E-P average distance and contour length, i.e., 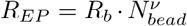, where *R_b_* denotes the scaling constant and v is the scaling exponent. To derive the scaling exponent defined by *ν* = [log(*R*) – log(*R_b_*)]/ log(*N*) we need to know the bead number N that corresponds to the real E-P DNA region. As a self-flexible chain, DNA on a small scale is rigid and tends to have a longer EED than an ideal flexible chain in a protein-free *in vitro* environment [23]. However, *in vivo*, as DNA is wrapped around nucleosomes, the DNA contour length in one (core) nucleosome (*L_nc_*) is around 50 nm, while the size of the (core) nucleosome (*R_nc_*) is around 10 nm [24]. Nucleosomes are linked by unfolded DNA. The linker length (*L_linker_*) is variable (0~80 bp) [24], and the linker’s EED can be approximated by *R_linker_* ≈ *L_linker_* owing to the rigidity of short DNA. We consider one bead composed of one (core) nucleosome and one linker. Thus, the DNA length in one bead *L_bead_* = *L_nc_* + *L_linker_*, and the size of one bead *R_bead_* ≈ *R_nc_* + *R_linker_*. Furthermore, the number of beads between enhancer and promoter

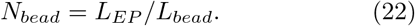

Finally, the average E-P distance can be expressed as

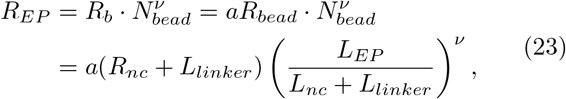

where *a* is constant, while *v* and *L_linker_* depend on the specific E-P experimental system. The walking spatial freedom of the E-P region could be characterized by deriving the scaling exponent *v* with Eq.23. However, the available experimental data are usually limited to determining *v*, *L_linker_*, and the scaling constant *a* together. For example, references [12] and [13] involve only two distinguishable lengths of E-P contour: short (3-5 *μ*m) and long (40-50 *μ*m). To derive *v* with these limited distinguishable contour lengths, we made the approximation *a* ≈ 1, i.e., *R_b_* ≈ *R_bead_*. The linker length is estimated to be 0 (for [13]) and 3 nm (for [12]), respectively, by letting *v* = 0.6 (3-D walk is assumed) for the long-contour E-P. The scaling exponent for the short-contour E-P can be derived around 0.8 (close to the 2-D walk exponent), which suggests that the short E-P DNA region may approximately walk in 2-D space. The high scaling exponent also excludes the possibility of enhancer and promoter walking inside the condensate.

By combining Eqs. 6 and 23, we can derive a theoretical estimate of the original distribution of E-P distance, i.e.,

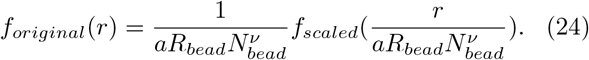

where *f_scaled_*(*r*) and *N_bead_* are defined by Eqs. 5 and 22, respectively. If we directly consider that the short E-P region walks in 2-D space, while the long E-P region walks in 3-D space, then the parameters *a* and *v* for a long or short contour, respectively, can be used in the same way as those for 3-D or 2-D space (see Eqs. 2 and 4 and Fig.7). As shown in Figure 5 (a) and Appendix Fig. 11 (a) and (c), the theoretically estimated 2-D or 3-D EED distribution is roughly consistent with the experimental distribution of the short or long E-P construct. This establishes the reasonableness of our hypothesis. The results could be understood based on the model in which enhancer communicates with promoter on the surface of a sufficiently large transcriptional condensate. Thus, when the E-P contour is short, TF binding sites are relatively enriched in the E-P region (i.e., a high density of confined sites) such that most of the E-P region is located on the surface of transcriptional condensate. However, when the E-P contour is long, TF binding sites are relatively sparse such that most of the E-P region walks outside of the transcriptional condensate. Moreover, 2-D-like walking space indicates the impossibility of the enhancer communicating with the promoter inside the condensate.

### E. Enhancer may dynamically activate promoter along the surface of transcriptional condensate

Naturally, one would wonder how the random walk of enhancers and promoters on the condensate surfaces affects transcriptional bursts. Here we propose a gene regulation model with the hypothesis that enhancer and promoter communicate on the surface of condensate. First, enhancers, promoters, and DNA fragments between them randomly walk on the transcriptional condensate surface. Second, when E-P distance is smaller than a threshold, the enhancer passes an activation signal to the promoter. Third, promoters can obtain a license to initiate the transcription independently and randomly (a Poisson process with licensing rate *λ_on_*). Fourth, the licensed promoter that is not activated by enhancer has a basic transcription level, which is not considered to be at the bursting state. Fifth, when both the license and activation signal are available at the promoter, then the transcription bursting occurs, corresponding to a higher transcription level. Sixth, losses of license and activation factors from the promoter are independent and random (Poisson) processes (loss rates λ_*off*,1_ and λ_*off*,2_). Seventh, transcription burst is delayed relative to the full activation of the promoter as time is always taken to allow RNA to accumulate to an experimentally detectable level [4, 13]. Based on the independence and stochasticity assumptions on the gain and loss of promoter licensing and enhancer activation signals, one can analytically derive the distribution of time difference between promoter licensing and enhancer activation for bursts as 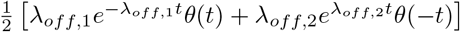 where *θ*(*t*) is a unit step function, and the distribution of burst duration as λ*e*^−λ*t*^, where λ = λ_*off*,1_ + λ_*off*,2_. With experimental timing profile of E-P distance and transcriptional bursts [12], we define promoter licensing time as that time when the transcription signal is detectable and the bursting time as that time when the transcription level is higher than a threshold. Enhancer activation time is that time when E-P distance is smaller than a threshold. With fitted λ_*off*,1_ and λ_*off*,2_, the simple theoretical model basically captures the distribution of the relative time obtained from experimental data [Fig. 6 (b)]. The predicted probability distribution of the duration of one transcription burst is also in good agreement with the experimental results [Fig. 6 (c)]. Therefore, our assumptions are reasonable, and based on these assumptions, we simulated E-P distance-dependent transcriptional bursts. The simulated probability distributions of the time of enhancer activation relative to promoter licensing and bursting duration are both consistent with the experimental and theoretical. If the delay of transcription burst relative to the promoter’s full activation is set to zero (i.e., the delay effect is removed), the simulation predicts that E-P distance during bursting is significantly smaller than that during non-bursting [Fig. 6 (d) and (e)], which conflicts with the slight difference in the distributions of experimental E-P distance [Fig. 12 (d)]. When the delay is randomly sampled from an exponential distribution with the average of 10 min, the simulation gives small differences [Fig. 6 (d) and (e)], similar to those of the experimental [Fig. 12 (c) and (d)]. This supports the existence of the delay effect in the real experimental system.

**FIG. 6.**
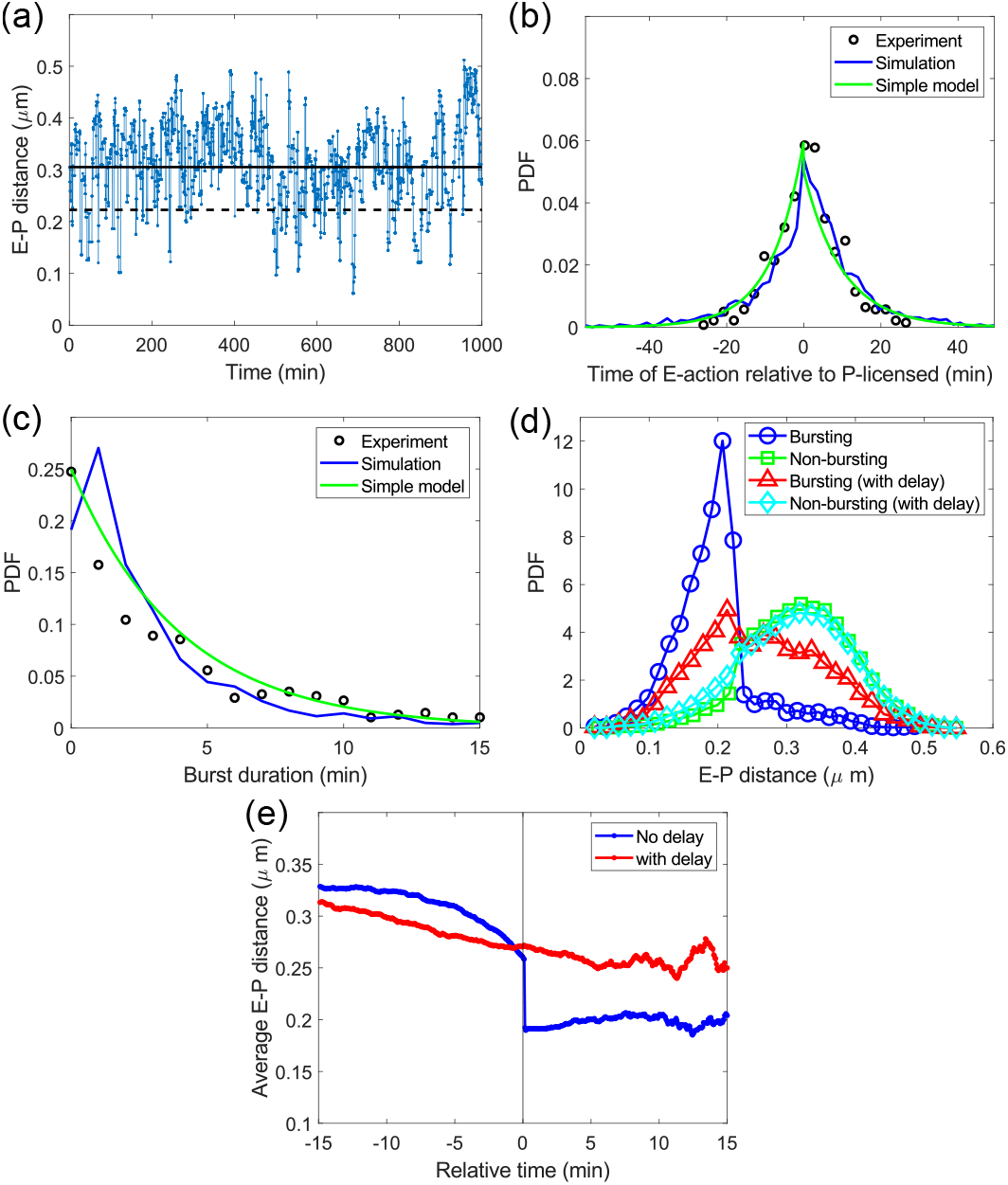
The simulated transcriptional bursts are comparable to those of experimental data [12]. (a) A profile of E-P distance changing with time which was arbitrarily chosen from the simulation. The black line indicates the average value, while the dashed black line indicates the average of E-P distances minus the standard deviation which was set as the threshold under which enhancer communicates with promoter. (b) The probability density function of the time of enhancer taking action on promoter (E-action) relative to the promoter obtaining the licensing signal (P-licensed) from both simulation and simple model is in agreement with the experimental. Simulation and simple model are based on the hypothesis of two independent Poisson processes. (c) The probability density function of transcriptional burst duration from simulation, along with the simple model, agrees with the experimental. (d) The probability distribution of E-P distance during bursting, or not bursting, with or without a fixed delay (10 min) of the transcription signal relative to full activation of the promoter. E-P distance tends to be smaller during bursting than during not bursting. However, the difference basically disappears with the delay. (e) Average E-P distance changes with time relative to start time of transcriptional burst. The average E-P distance has a fast decrease at the burst start time, but the decrease slows down if a delay is added to the transcription burst after licensing and activating the promoter. Details of determining the parameters are given in Appendix D.

**FIG. 7.**
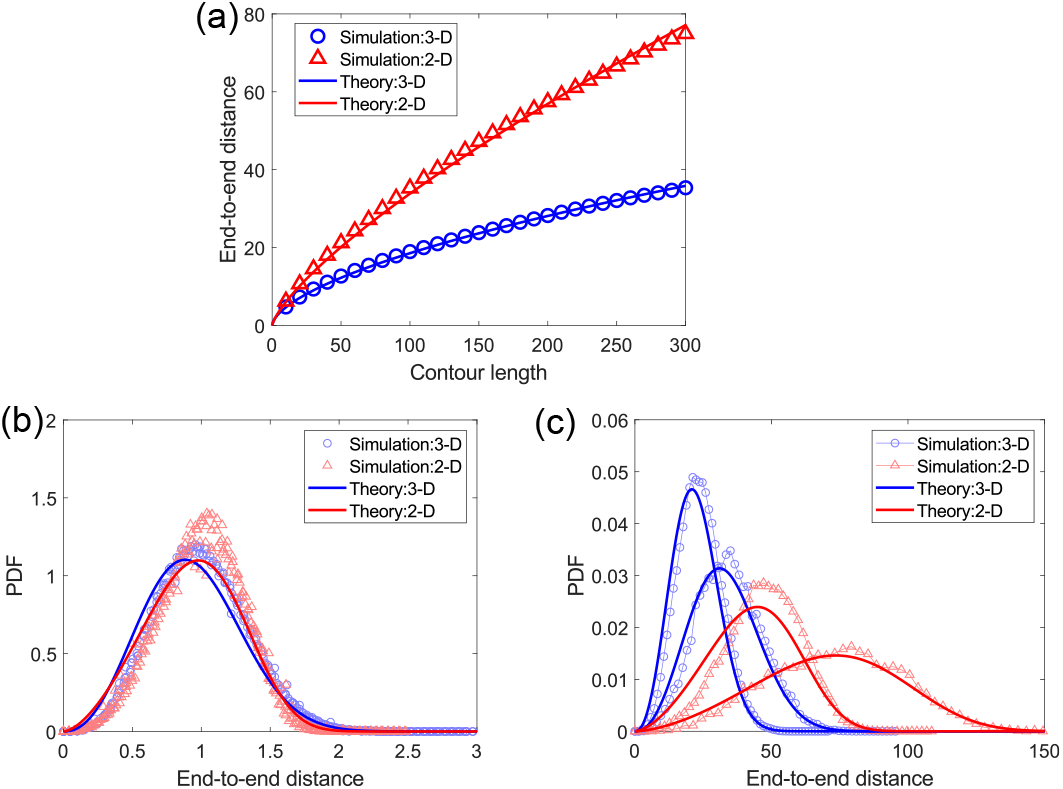
Classical theories properly characterize the simulated random walk of a self-avoiding chain (bead number=401) in 3-D or 2-D Euclidean space. (a) EED is a power function of contour length as predicted by the Flory theory. By fitting the simulation with Eq. 2 or 4, the constant *a* was determined to be 1.17 (3-D) or 1.07 (2-D). (b) The distribution of scaled EED (divided by the average) nearly collapses together and agrees with theoretical predictions. (c) The distribution of EED can be estimated by combining Flory theory and theoretical formulas for scaled distribution.

**FIG. 8.**
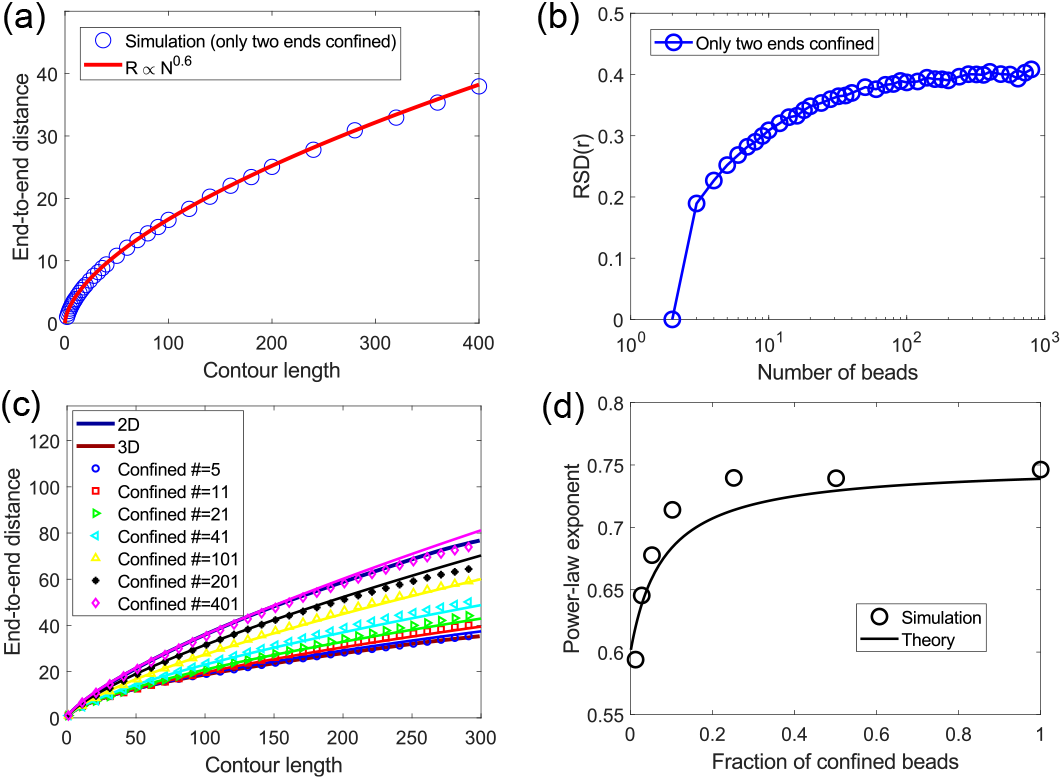
The scaling theory characterizes the random walk of DNA chain with some beads confined to the surface of a large sphere (*R_S_* = 10^4^). (a) The simulated chain with only two ends confined to the spherical surface, such that contour length between the two confined ends was changed, shows that EED increases along with contour length in a power-law relationship *R* ∝ *N*^0.6^. (b) Relative standard deviation (RSD) of EED *r* increases with contour length (bead number).(c) The relationship between EED and contour length is changed according to the number of confined beads. Confined beads few in number lead to the 3D result, whereas confinement of all beads leads to the 2D result. Symbols (different confined numbers) and thick lines (2D and 3D) denote results from simulation; other lines denote theoretical results obtained by fitting simulation results with Eq.14. Parameters: *b* = 1.2 and *g_c_* = 12.5. (d) The scaling method shows that the powerlaw exponent increases with the proportion of confined beads from ~0.6 (3D exponent) to ~0.75 (2D exponent) in rough agreement with the simulation.

**FIG. 9.**
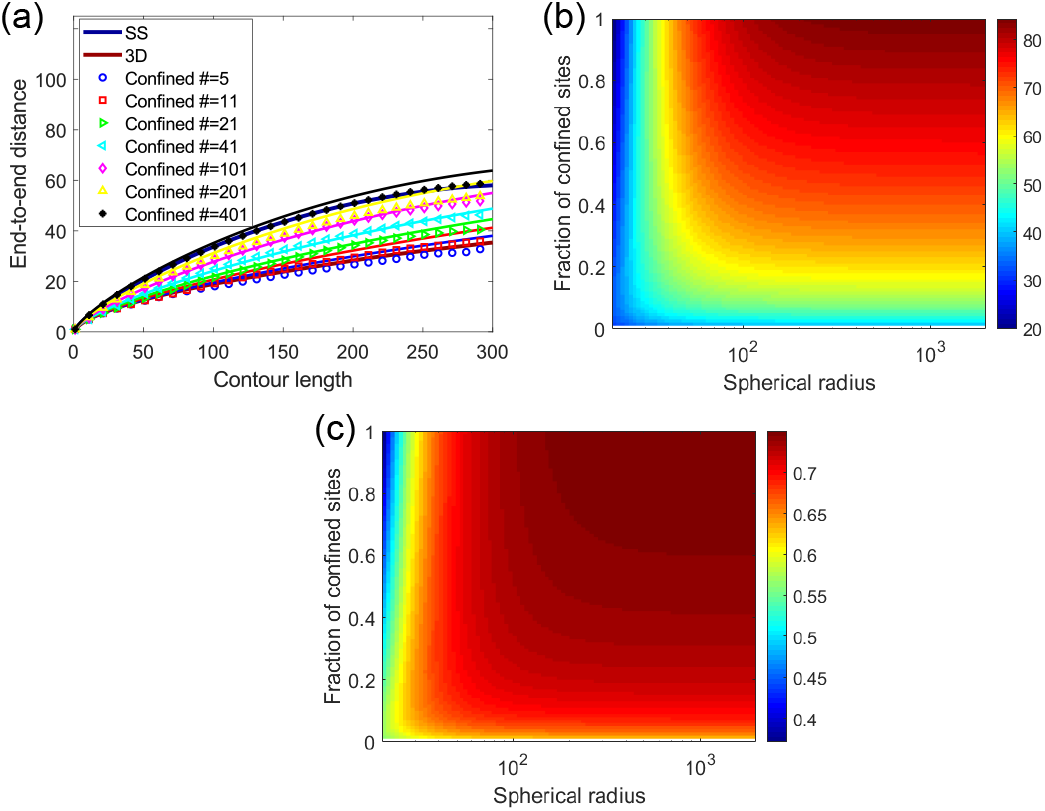
The dependence of EED and power-law exponent on both spherical size and confined bead proportion when the chain walks on the sphere. The relevant parameters used here are the same as those for Figures 1 and 2. (a) EED as a function of contour length decreases along with the number of beads confined to the surface of a small sphere (*R* = 40). (b) EED as a function of spherical size and confined proportion given contour length (=300). (c) Power-law exponent as a function of spherical size and confined proportion.

**FIG. 10.**
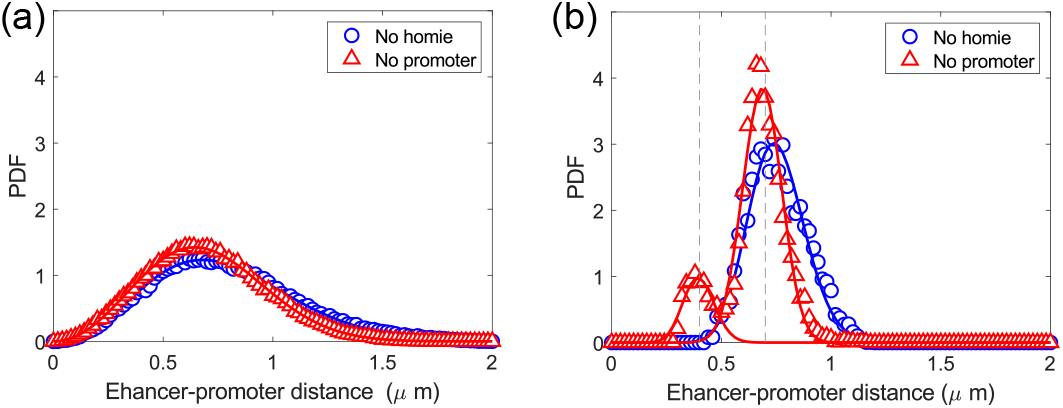
The separation of the paired and unpaired states for constructs with homie insulators [12] by smoothing the E-P distance timing profile. The function *Ar^g^* exp(-*αr^δ^*) was used to fit the probability distribution of E-P distance (*r*). The green line represents the construct without promoter (no promoter), while the blue line represents the construct without homie insulator (no homie insulator). (a) One peak exists in the probability distribution of E-P distance for the construct without homie insulator or promoter. Fitting parameters: *A* = 8.36, *g* = 2.05, *α* = 2.45, and *δ* = 2.36. (b) Two peaks emerge from the probability distribution of smoothed E-P distance for the construct without promoter; however, there is still only one peak for the construct without homie insulator. The E-P distance timing profile was smoothed with a sliding window of 15 min. The fitting for no homie was done for E-P distance longer than 0.5 *μm*. The fitting for no promoter was done for E-P distance longer and shorter than 0.5 *μm* separately. Dashed gray lines show the distance thresholds at 0.4 and 0.7 *μm*, respectively. Parameters: *A* = 2.86 × 10^7^, *g* = 18.7, *α* = 17.92, *δ* = 1.79 (no homie); *A* = 4.9 × 10^5^, *g* = 17.48, *α* = 18.39, *δ* = 3.37 (no promoter, E-P distance> 0.5*μm*); *A* = 1.67 × 10^8^, *g* = 13.63, *α* = 48.09, *δ* = 2.19 (no promoter, E-P distance< 0.5*μm*).

**FIG. 11.**
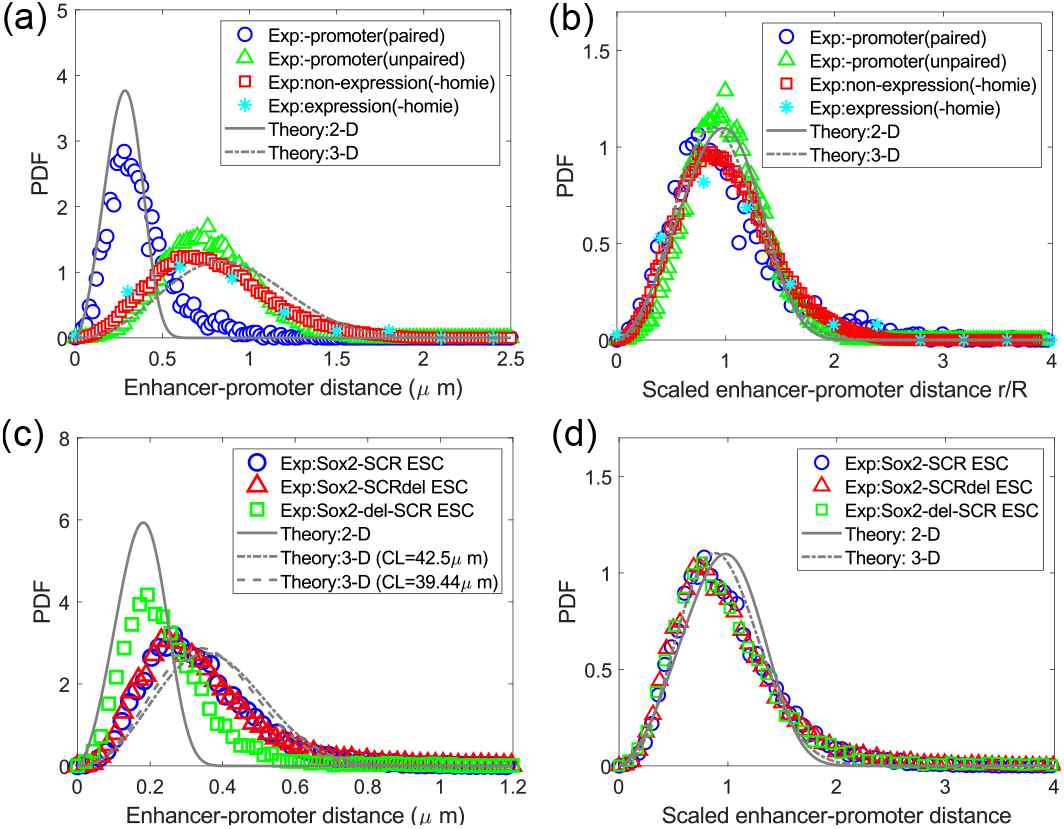
Probability distribution of E-P distance with different contour lengths is captured by the Flory theory and scaling method. Experimental data from Chen et al. [12] and Alexander et al. [13] were used. Parameters are given in Appendix C2. (a,c) Probability density function (PDF) of original E-P distance was well estimated by the theory described by Eq. 24. (b,d) Probability density function (PDF) of scaled E-P distance collapses to the theoretical 2-D and 3-D scaled distributions when E-P distance is divided by its average.

**FIG. 12.**
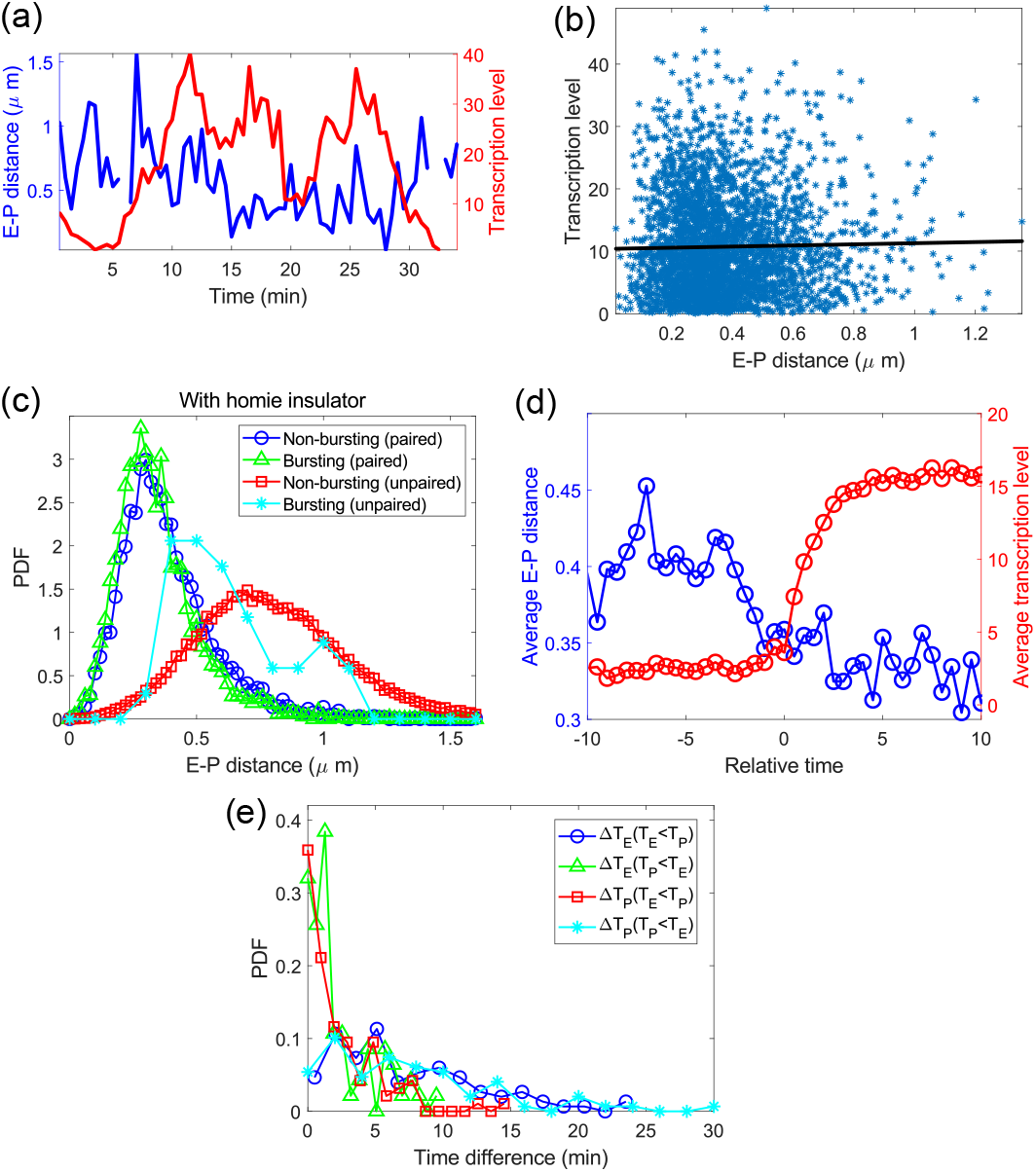
A weak correlation between E-P distance and transcription shown by single-molecule experiments [12]. (a) Profiles of E-P distance and transcriptional level changing with time. (b) Scatter plot of transcriptional level versus E-P distance. Black line represents the linear fit. Pearson correlation coefficient is 0.02; p-value = 0.27. (c) Probability distribution of E-P distance during bursting or non-bursting when the homie insulator is paired or unpaired. (d) Average E-P distance and average expression level change with the relative time contrarily. The profiles during paired periods were aligned according to the time when the burst occurs. (e) Probability distribution of the time difference between *T_E_* (or *T_P_*) and *T_EP_*, when *T_P_* < *T_E_* or *T_E_* <*T_P_*.

## IV. DISCUSSION

In the present work, we proposed a novel model for EP communication on the surface of transcriptional condensate. The model was evidenced by comparison with experimental data on several characteristic walking parameters or variables, e.g., the power-law exponent in the relationship between average EED and contour length and the distribution of EED. Under many spatial limitations, the EED of a polymer chain (linear connection of beads) is approximately a power function of contour length (bead number *N*). The power-law exponent *ν* is a key parameter that reflects the spatial degree of freedom of random walk of the polymer chain. Moreover, EED distribution is featured by the scaling law since it collapses after dividing by *N^v^*. Because of the limitation of distinguishable E-P contour lengths in the usable experimental data, the scaling exponent *ν* was estimated by (log *R_EP_* – log *R_bead_*)/log *N*, rather than (log *R_EP_* – log *R_b_*)/log *N*, at a given bead number *N* (see Eq.23). The scaling constant *R_b_*/*R_bead_* is usually a bit greater than 1, according to the above results. The small deviation of *R_b_/R_bead_* from 1 would not affect the scaling exponent much. Therefore, the less exact exponent still could roughly characterize the spatial freedom of random walking of the E-P DNA region. However, a more exact scaling exponent is still required to further verify our results. Therefore, special single-molecule tracking experiments with more than two distinguishable E-P contour lengths need to be done.

The E-P interaction mechanisms studied here also reflect featured location distributions of DNA around transcriptional condensates. The communication of enhancer and promoter along the condensate surface or interior requires confinement of DNA sites to the condensate surface or inside the condensate. The confinement of DNA sites can be mainly achieved by specific binding with TFs. Thus, an issue is raised about the location of TFs in transcriptional condensates. If the binding of the confined beads to TFs is stronger than the repulsion of the unconfined beads to the condensate, then the DNA chain may frequently enter the condensate by TF pulling. If the repulsion of the unconfined beads to the condensate is stronger than the binding of the confined beads to TFs, then the E-P DNA region, as well as bound TFs, may stay near the surface of the condensate. If DNA-bound TFs occupy a high ratio among all TFs inside the condensate, one would observe an enrichment of TFs on the condensate surface. More advanced single-molecule detection techniques are required to precisely locate the E-P DNA region and bound TFs.

The dynamic kissing mechanism of E-P communication is built on the belief that transcription activation requires close proximity between enhancer and promoter. Chen et al. have found that E-P distance is reduced when transcription is on, compared to when transcription is off [12]. However, other researchers found that E-P proximity is not necessary for transcription activation [4, 10, 13]. In the experimental E-P system used by Chen et al. [12], a pair of insulators (homies) is involved, and the pairing of homies makes the effective contour length of E-P much smaller. The transcription on and off states can be further partitioned into homie-paired and -unpaired states. Analyses show that E-P distance still decreases at on state compared with off state, no matter whether the homies are paired or unpaired, but the decreases are small (see [12] and Appendix Figure 12). The observed little, or weak, correlation between EP distance and transcription may result from the inability of delayed detection of transcription signals to keep up with the dynamic communication between enhancer and promoter [4, 13]. Thus, new measurement techniques cancelling the delay are needed to further establish the correlation.

In our gene regulation model, we assumed that the occurrence of transcription bursts requires both the initiation license of the promoter and the activation of enhancer. Several candidate mechanisms may serve to provide such license and the signal that the enhancer transmits to the promoter. For example, it is known that phosphorylation in the C-terminal domain of preinitiation complex of transcription and other core proteins is required for transcription initiation [25, 26] and that enhancers may affect the phosphorylation of those domains by recruiting necessary kinases [26].

Through our integrated framework, we propose an imaginative model of long-range E-P communication. First, enhancer and promoter were roughly drawn together by reducing the effective contour length between them. This process might be governed by looping mechanisms that could be mediated by flanking factors [4, 27, 28]. Second, formation of transcriptional condensate that may be driven by liquid-liquid phase separation could be prevalent in the nucleus [4, 10]. Third, both enhancer and promoter may be maintained on the surface of transcriptional condensate, which may be caused by the balance between condensate-DNA repulsion and DNA binding TFs-condensate attraction. Fourth, enhancers randomly approach and activate promoters along the surface of transcriptional condensate. It is worth noting that our model does not completely exclude the alternative dynamic kissing models [4, 29], or looping model [4, 27, 28]. Nevertheless, the analytic and empirical results presented here provide a solid basis for the E-P communication-on surface model.

## V. CONCLUSION

In this study, we applied the classic SAW model to simulate the random walk of E-P region under various spatial constraints. Combining the Flory theory and scaling method, which are classic in polymer physics, we characterized the relationship between average EED and contour length, as well as the distribution of EED, and their dependence on spatial constraint parameters. By comparing the walking features of a polymer chain, either fully or partially confined on spherical surface or interior, with classic spaces (2-D and 3-D), we excluded the possibility of enhancers and promoters walking inside a spherical condensate or on the surface of a small spherical condensate. Based on the results, we, for the first time, pointed out that both enhancer and promoter may walk on the surface of a large sphere, possibly achieved by binding of TFs with limited DNA sites and the repulsion of the condensate to DNA. Based on the timing profiles of E-P distance and transcription level, we further proposed a novel enhancer-mediated transcriptional regulation model whereby the enhancer dynamically kisses and activates the promoter along the transcriptional condensate surface. More experiments will be carried out to test this detailed mechanism.

## ACKNOWLEDGMENTS

This work was supported by the Beijing Natural Science Foundation (Z200021), Special Investigation on Science and Technology Basic Resources of MOST, China (2019FY100102), the Strategic Priority Research Program of the Chinese Academy of Sciences, China (X-DA24020307), the National Key R&D Program of China (2018YFC2000400) and the National Nature Science Foundation of China (91940304, 31871331,31671342). We thank Dr. Wulan Deng (Peking University) and her group for helpful discussion.

## Appendix A: Classical theories properly characterize the random walks of DNA chain in 3-D or 2-D Euclidean space

As shown in Figure 7 (a), the simulated EED as a function of contour length can be well captured by Eqs. 2 and 4. The probability distribution of the scaled EED (*r/R*) is in rough agreement with the theoretical formulas shown by Eq. 5, whether in 3-D or 2-D space (Fig. 7 (b)). Figure 7 (b) also shows that the 2-D and 3-D EED distributions nearly collapse together by scaling. Combining the formulas for estimating the average of EED and probability distribution of the scaled EED, we obtained the formula for probability distribution of original EED (Eq. 6), which basically captured the simulated original distribution [Fig. 7 (c)].

## Appendix B: The scaling theory can also characterize the random walk of DNA chain with some beads confined to a spherical surface

Usually, the scaling and Flory theories can both be used to characterize the random walk of a self-avoiding polymer chain [17]. The Flory theory is built on the balance of the free energy contributed by entropy and the interaction energy between chain beads and between the chain and spatial boundary. The scaling theory is based on the scaling law features of the polymer chain. These two theories can be regarded as two sides of the same coin because they link EED to contour length from different views. Flory theory gives a mechanistic result, while the scaling theory shows a universal and phenomenological result. Thus, it is understandable that they both (represented by Eqs. 11–15) well captured the average EED as a function of contour length and confined bead proportion when limited beads of the polymer chain are confined to a spherical surface [Fig. 2 (b), (c) and Fig. 8 (c)] and the increase in the power-law exponent according to the proportion of confined beads [Fig. 2 (f) and Fig. 8 (d)]. Equations 14 and 15 can also combine with Eq. 9 to estimate the mixed effects of condensate size and confinement proportion. The results are similar to those given by Eqs. 13 and 16 and shown by Figure 9.

## Appendix C: Details of analyzing single-molecule tracking data

Some details of analyzing experimental data from literature, including references [12] and [13] are supplemented below.

### 1. Separation of paired and unpaired states for constructs with homie insulator in reference [12]

Most experimental constructs of Chen et al. [12] include two homie insulators which can be paired to form a loop. In principle, E-P distance is shorter when homie insulators are paired than when homie insulators are unpaired. However, only one peak is observed in the probability distribution of E-P distance, e.g., for the construct without promoter, as shown in Figure 10 (a). Considering the continuity of the paired state, we used a sliding window (15 min) to smooth the E-P distance timing profile, taking the average within the window. The smoothed probability distribution of E-P distance has two well-separated peaks [Fig. 10 (b)]. The paired state should mainly contribute to the smaller left peak, while the unpaired state should mainly contribute to the bigger right peak. As a control, the distribution for the construct without homie insulator does not show two peaks, irrespective of smoothing (Fig. 10). Based on the smoothed probability distribution, we set two distance thresholds (0.4 *μm* and 0.7 *μm*) to isolate the paired and unpaired states. When the average E-P distance is shorter than the smaller threshold, the system is considered at the paired state. When the average distance is longer than the greater threshold, the system is at the unpaired state. The smaller threshold we chose is closer to the paired peak, while the greater threshold is closer to the unpaired peak because the two states are likely to be mixed in the intermediate range between two thresholds. Given the paired and unpaired states, further analyses on the original E-P distance timing profile were performed. Especially, the contour length between enhancer and promoter at the unpaired state is defined as their sole distance along the non-intersecting E-P DNA region, while the effective contour length at the paired state is defined as their closest distance along the genome cross structure formed by homie insulator paring.

### 2. The parameters for determining the power-law exponent with experimental data

Four groups of experimental data from [12] were used for Figure 5 based on whether insulators (homies) are paired or unpaired and if the gene is transcribing (expression) or not (non-expression): (I) non-expression (paired), (II) expression (paired), (III) non-expression (unpaired), and (IV) expression (unpaired). Parameters for Figure 5: E-P contour length for unpaired insulators is 52 *μm*; effective contour length for paired insulators is 3.06 *μm*; *L_linker_* = 3 nm, *L_nc_* = 50 nm, *d_nc_* = 10 nm. The average E-P distances for groups (I)-(IV) are 0.4, 0.35, 0.79, and 0.68 *μm*, respectively. The derived power-law exponent is 0.84, 0.81, 0.6, and 0.58, respectively.

Four groups of experimental data from [12] were used for Figure 11 (a) and (b) based on whether 1) the promoter or homie insulator is knocked out (-promoter or -homie), 2) the insulators (homies) are paired or unpaired, and 3) the gene is transcribing (expression) or not (non-expression): (I) -promoter(paired), (II) - promoter(unpaired), (III) non-expression(-homie), and (IV) expression(-homie). Parameters for Figure 11 (a) and (b): E-P contour length= 52 (unpaired insulators) or 3.06 *μm* (paired insulators); *d_linker_* = *L_linker_* = 3 nm, *L_nc_* = 50 nm, *d_nc_* = 10 nm. The average E-P distances for datasets (I)-(IV) are 0.37, 0.76, 0.78, and 0.75 *μm*, respectively. The derived power-law exponent is 0.83, 0.59, 0.59, and 0.59, respectively.

The single-molecule tracking data of three constructs from [13] were used for Fig.11 (c) and (d): (I) Sox2-SCR ESC, (II) Sox2-SCRdel ESC, and (III) Sox2-del-SCR ESC. Parameters for Fig.11 (c) and (d): contour length = 42.5 (I), 39.44 (II), or 4.42 *μm* (III); *d_linker_* = *L_linker_* = 0 nm, *L_nc_* = 50 nm, *d_nc_* = 6 nm. The average E-P distances for datasets (I)-(III) are 0.34, 0.34, and 0.25 *μm*, respectively. The derived power-law exponents are 0.6, 0.6, and 0.83, respectively.

### 3. Correlation between E-P distance and transcription level

From the timing profile and scatter plot [Fig.12 (a) and (b)], no significant correlation was observed between E-P distance and transcription level. The Pearson correlation coefficient between E-P distance and transcription level is small (0.02), and the corresponding p-value is high (0.27). As mentioned in the main text, the transcription signal may be delayed relative to E-P communication. Accordingly, we also computed the correlation between E-P distance and transcription level with a pre-given constant delay, but we still did not obtain any significant correlations. This could be explained by the delay of transcription level which is a random variable owing to the randomness of the transcription process. We defined the burst periods as occuring when the transcription level is higher than a threshold (typically, 5). From the data for the species with insulator, the distribution of E-P distance during bursting also deviates from that during non-bursting. That is, at the paired state, the deviation is small, while at the unpaired state, the deviation is a little greater [Fig.12 (c)]. We aligned the timing profiles during paired periods according to the time when the burst occurs. The obtained average expression level shows a sharp increase after the burst begins, while the average E-P distance tends to decrease before the burst [Fig.12 (d)]. These results show a weak correlation between E-P distance and transcription level.

### 4. Timing sequences of promoter licensed and enhanced

As we assumed in the main text, the promoter is licensed by some signals and enhanced by enhancer independently. A non-licensed promoter cannot initiate transcription, licensed, but non-enhanced promoter, can initiate transcription at a low level, while licensed and enhanced promoter can initiate transcription at a high level. We learned timing sequences of promoter licensed and enhanced from timing profiles of E-P distance and transcription level. The time of promoter licensed (*T_P_*) occurs at the start of the expression period, while the time of promoter enhanced (*T_E_*) occurs when E-P distance becomes short enough to reach the corresponding threshold, typically the average of E-P distance minus its standard deviation. The time of full promoter activation (*T_EP_*) occurs when the transcription level is higher than the corresponding threshold, typically 5. In principle, *T_EP_* should be equal to max(*T_P_,T_E_*) for one burst. To test this result, we computed probability distributions of Δ*T_E_* = *T_EP_* – *T_E_* and Δ*T_P_* = *T_EP_* – *T_P_*. As expected, when *T_P_* > *T_E_*, Δ*T_P_* has a narrow distribution close to zero, while Δ*T_E_* has a wide distribution. When *T_P_* < *T_E_*, Δ*T_E_* has a narrow distribution close to zero, while Δ*T_P_* has a wide distribution [Fig. 12 f].

## Appendix D: Simulation of transcription bursting

Based on assumptions made in the main text, we simulated transcription bursting. Here we present the determination of the parameters involved: First, loss rates of the promoter’s license and activation signals, respectively λ_*off*,1_ and λ_*off*,2_), were fixed by fitting the experimental time difference between promoter licensed and enhancer activation with the following function: 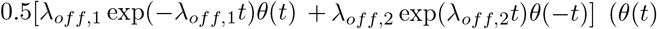 is a unit step function); then λ_*off*,1_ = 0.11 min^−1^, and λ_*off*,2_ = 0.14min^−1^. The time interval between two adjacent simulation steps (i.e., the time for one walk of the chain) was determined by fitting the simulated burst duration with the experimental. The fitted step interval is 0.05 min. The promoter-licensed rate is arbitrarily chosen: λ_*on*_ = 0.15 min^−1^. The usage of a lower (0.015 min^−1^) or higher λ_*on*_ (1.5 min^−1^) does not affect our main results much, but it did change the bursting frequency significantly. Based on the independence of promoter licensing and enhancer activation, the probability distribution of burst duration can be estimated by the simple function λexp(–λ*t*), where λ = λ_*off*,1_ + λ_*off*,2_. This function well captures the burst duration distribution, as shown in Figure 6 (c). The threshold of E-P communication distance was defined as the average of E-P distance (〈*r*〉) minus its standard deviation (*σ*(*r*)). We also tested lower and higher thresholds (e.g., 〈*r*〉 – 1.5*σ*(*r*) and 〈*r*〉 – 0.5*σ*(*r*)), which did not substantially change our results.

## Notes

### Competing Interest Statement

The authors have declared no competing interest.

## References

[1] D. Hnisz, B. J. Abraham, T. I. Lee, A. Lau, V. Saint-André, A. A. Sigova, H. A. Hoke, and R. A. Young, Super-enhancers in the control of cell identity and disease, Cell 155, 934 (2013).

[2] M. Levine, C. Cattoglio, and R. Tjian, Looping back to leap forward: transcription enters a new era, Cell 157, 13 (2014).

[3] B. Lim, T. Heist, M. Levine, and T. Fukaya, Visualization of transvection in living drosophila embryos, Molecular cell 70, 287 (2018).

[4] H. B. Brandão, M. Gabriele, and A. S. Hansen, Tracking and interpreting long-range chromatin interactions with super-resolution live-cell imaging, Current Opinion in Cell Biology 70, 18 (2021).

[5] A. Raj, C. S. Peskin, D. Tranchina, D. Y. Vargas, and S. Tyagi, Stochastic mrna synthesis in mammalian cells, PLoS biology 4, e309 (2006).

[6] D. M. Suter, N. Molina, D. Gatfield, K. Schneider, U. Schibler, and F. Naef, Mammalian genes are transcribed with widely different bursting kinetics, science 332, 472 (2011).

[7] B. R. Sabari, A. Dall’Agnese, A. Boija, I. A. Klein, E. L. Coffey, K. Shrinivas, B. J. Abraham, N. M. Hannett, A. V. Zamudio, J. C. Manteiga, C. H. Li, Y. E. Guo, D. S. Day, J. Schuijers, E. Vasile, S. Malik, D. Hnisz, T. I. Lee, I. I. Cisse, R. G. Roeder, P. A. Sharp, A. K. Chakraborty, and R. A. Young, Coactivator condensation at super-enhancers links phase separation and gene control, Science 361 (2018).

[8] A. Boija, I. A. Klein, B. R. Sabari, A. Dall’Agnese, E. L. Coffey, A. V. Zamudio, C. H. Li, K. Shrinivas, J. C. Manteiga, N. M. Hannett, et al., Transcription factors activate genes through the phase-separation capacity of their activation domains, Cell 175, 1842 (2018).

[9] D. Hnisz, K. Shrinivas, R. A. Young, A. K. Chakraborty, and P. A. Sharp, A phase separation model for transcriptional control, Cell 169, 13 (2017).

[10] B. Lim and M. S. Levine, Enhancer-promoter communication: hubs or loops?, Current Opinion in Genetics & Development 67, 5 (2021).

[11] N. S. Benabdallah, I. Williamson, R. S. Illingworth, L. Kane, S. Boyle, D. Sengupta, G. R. Grimes, P. Therizols, and W. A. Bickmore, Decreased enhancer-promoter proximity accompanying enhancer activation, Molecular cell 76, 473 (2019).

[12] H. Chen, M. Levo, L. Barinov, M. Fujioka, J. B. Jaynes, and T. Gregor, Dynamic interplay between enhancer–promoter topology and gene activity, Nature genetics 50, 1296 (2018).

[13] J. M. Alexander, J. Guan, B. Li, L. Maliskova, M. Song, Y. Shen, B. Huang, S. Lomvardas, and O. D. Weiner, Live-cell imaging reveals enhancer-dependent sox2 transcription in the absence of enhancer proximity, elife 8, e41769 (2019).

[14] Y. Shin, Y.-C. Chang, D. S. Lee, J. Berry, D. W. Sanders, P. Ronceray, N. S. Wingreen, M. Haataja, and C. P. Brangwynne, Liquid nuclear condensates mechanically sense and restructure the genome, Cell 175, 1481 (2018).

[15] D. S. Lee, N. S. Wingreen, and C. P. Brangwynne, Chromatin mechanics dictates subdiffusion and coarsening dynamics of embedded condensates, Nature Physics 17, 531 (2021).

[16] Y. Zhang, D. S. Lee, Y. Meir, C. P. Brangwynne, and N. S. Wingreen, Mechanical frustration of phase separation in the cell nucleus by chromatin, Physical review letters 126, 258102 (2021).

[17] M. Rubinstein, R. H. Colby, et al., Polymer physics, Vol. 23 (Oxford university press New York, 2003).

[18] N. Madras and A. D. Sokal, The pivot algorithm: a highly efficient monte carlo method for the self-avoiding walk, Journal of Statistical Physics 50, 109 (1988).

[19] D. McKenzie and M. Moore, Shape of self-avoiding walk or polymer chain, Journal of Physics A: General Physics 4, L82 (1971).

[20] J. Des Cloizeaux, Lagrangian theory for a self-avoiding random chain, Physical Review A 10, 1665 (1974).

[21] S. Caracciolo, M. S. Causo, and A. Pelissetto, End-to-end distribution function for dilute polymers, The Journal of Chemical Physics 112, 7693 (2000).

[22] K. Shrinivas, B. R. Sabari, E. L. Coffey, I. A. Klein, A. Boija, A. V. Zamudio, J. Schuijers, N. M. Hannett, P. A. Sharp, R. A. Young, et al., Enhancer features that drive formation of transcriptional condensates, Molecular cell 75, 549 (2019).

[23] F. Valle, M. Favre, P. De Los Rios, A. Rosa, and G. Dietler, Scaling exponents and probability distributions of dna end-to-end distance, Physical review letters 95, 158105 (2005).

[24] J. Widom, Toward a unified model of chromatin folding, Annual review of biophysics and biophysical chemistry 18, 365 (1989).

[25] D. Nikolov and S. Burley, Rna polymerase ii transcription initiation: a structural view, Proceedings of the National Academy of Sciences 94, 15 (1997).

[26] C.-T. Ong and V. G. Corces, Enhancer function: new insights into the regulation of tissue-specific gene expression, Nature Reviews Genetics 12, 283 (2011).

[27] W. Deng, J. Lee, H. Wang, J. Miller, A. Reik, P. D. Gregory, A. Dean, and G. A. Blobel, Controlling long-range genomic interactions at a native locus by targeted tethering of a looping factor, Cell 149, 1233 (2012).

[28] M. Merkenschlager and E. P. Nora, Ctcf and cohesin in genome folding and transcriptional gene regulation, Annual review of genomics and human genetics 17, 17 (2016).

[29] W.-K. Cho, J.-H. Spille, M. Hecht, C. Lee, C. Li, V. Grube, and I. I. Cisse, Mediator and rna polymerase ii clusters associate in transcription-dependent condensates, Science 361, 412 (2018).

